# An Exploratory Study of Prefrontal Cortex Activation Related to Golf Putting Performance Under Psychological Pressure: A Functional Near-infrared Spectroscopy Approach

**DOI:** 10.64898/2026.07.13.737385

**Authors:** Ryosuke Hiyama, Norito Nakata, Ren Inoue, Takahiro Manai, Tatsuo Hirata, Hiroki Sato

**Author notes:** Corresponding Author: Hiroki Sato^1, 2^ 307 Fukasaku, Minuma-ku, Saitama 337-8570, Japan.

## Abstract

Neurofeedback (NF) is a promising method for helping individuals overcome choking under pressure. Identifying relevant neural biomarkers is crucial for developing effective NF. In this exploratory study, we aimed to investigate changes in prefrontal hemodynamic signals associated with golf-putting performance under psychological pressure using functional near-infrared spectroscopy. Participants engaged in a one-on-one golf-putting task against an experimenter, with monetary rewards introduced to induce psychological pressure. This manipulation successfully elicited psychological pressure, leading to impaired performance in some participants. Based on performance changes between the practice and competition sessions, participants were categorized into a *non-choking group* (performance improved) and a *choking group* (performance declined). Statistical analysis revealed significantly greater increases in prefrontal activation from practice to competition in the non-choking group than in the choking group, especially in the left superior frontal gyrus. Furthermore, moderate but statistically nonsignificant negative correlations were observed between changes in activation in this region and changes in putting error, indicating that greater activation increases tended to accompany less performance deterioration or greater performance improvement. These exploratory findings suggest that the left superior frontal gyrus warrants further investigation as a candidate biomarker for NF interventions aimed at mitigating choking under pressure.

## Introduction

In situations involving psychological pressure, such as important sports competitions or musical performances, performing at one’s usual level of skill or replicating daily practice outcomes is often challenging. Factors such as the outcome of the game, the presence of spectators, and anxiety about failure can interfere with normal performance. Baumeister (1984) defined psychological pressure as “a combination of factors that increase the importance of achieving high performance in a specific situation” and described the resultant performance decline under such pressure as “choking under pressure” (hereafter referred to simply as *choking*). Subsequently, Mesagno and Hill (2013) offered a more comprehensive definition, describing choking as “an acute and considerable decrease in skill execution and performance, despite normally achievable self-expected standards, resulting from increased anxiety under perceived pressure” (Mesagno & Hill, 2013; Gröpel & Mesagno, 2019). This phenomenon is not limited to individuals with high levels of expertise; even laypersons often experience choking in their everyday lives.

Neurofeedback (NF) is a promising approach to overcoming choking (Kohl et al., 2020). NF uses various neuroimaging modalities, such as electroencephalography (Marzbani, Marateb & Mansourian, 2016), functional magnetic resonance imaging (fMRI) (Sorger et al., 2018), and functional near-infrared spectroscopy (fNIRS), to monitor neural and hemodynamic activity in real time (Kohl et al., 2020). Participants receive these signals as visual or auditory feedback, enabling them to learn how to regulate their brain activity. The ultimate goal is to achieve self-regulation without external feedback (Sorger et al., 2018). As NF provides direct feedback on the quantitative indices of mental states, it represents a promising strategy for mitigating choking.

A recent meta-analysis showed the feasibility of using NF to improve motor performance (Onagawa et al., 2023). These findings suggest that NF can positively influence motor performance through the self-regulation of brain activity, suggesting its potential use in specific situations, such as performance under pressure.

However, to develop effective NF systems, understanding brain activity associated with performance fluctuations under psychological pressure is essential, as this knowledge is crucial for identifying appropriate biomarkers. Previous studies have reported differences in brain activity between successful and unsuccessful performances under pressure (Cooke et al., 2014; Slutter, Thammasan & Poel, 2021). However, these studies were not conducted within the conceptual framework of choking as defined by Mesagno and Hill (2013); rather, they examined performance under pressure in a broader sense. Notably, not all individuals experience a decline in performance when under pressure; some may even improve (Hibbs, 2010). To date, no study has simultaneously examined performance changes and brain activity while accounting for individual differences, using each participant’s baseline performance as a reference.

Collectively, existing findings have advanced our understanding of the mechanisms underlying choking and its possible countermeasures; however, further investigations within a more rigorous theoretical framework are required. Therefore, we aimed to explore optimal targets for NF intervention. Specifically, we used a golf-putting task to establish the baseline performance for each participant and applied fNIRS to investigate brain activity under conditions in which performance declined relative to baseline due to psychological pressure compared with conditions in which performance was maintained. In addition, we aimed to explore the neural features that could serve as reliable biomarkers for future NF-based interventions aimed at overcoming choking.

## Materials and method

### Participants

Twenty male students (mean age 21.3 ± 1.3 years) participated in this study. The sample size was determined heuristically based on preliminary experiments (Lakens, 2022). The Edinburgh Handedness Inventory was used to confirm that all participants were right-handed (Oldfield, 1971) and inexperienced in golf. The Ethics Committee of Shibaura Institute of Technology approved the study protocol (Application No. 19-008, Toyosu Academic Affairs Division Document No. 19128), and all participants provided written informed consent prior to their participation.

### Task and procedure

Participants participated in a one-on-one putting competition with the experimenter across two sessions: a practice session and a competition session. In each session, they performed five trials of a 1.5-m putt (Smith et al., 2000; Balk et al., 2013). The experimenter always putted first, followed by the participant. The shortest distance between the final position of the ball and the center of the target cross was defined as the error value and used as an indicator of motor performance.

Each trial lasted 45 s, consisting of a 5-s rest period, a 10-s focus period devoted to putting, and a 30-s rest period. Experimental stimulus presentation software (Presentation, Neurobehavioral Systems) provided auditory cues at the beginning and end of the focus period. Participants were instructed to begin putting immediately after the second cue (Fig. 1).

**Figure 1.**
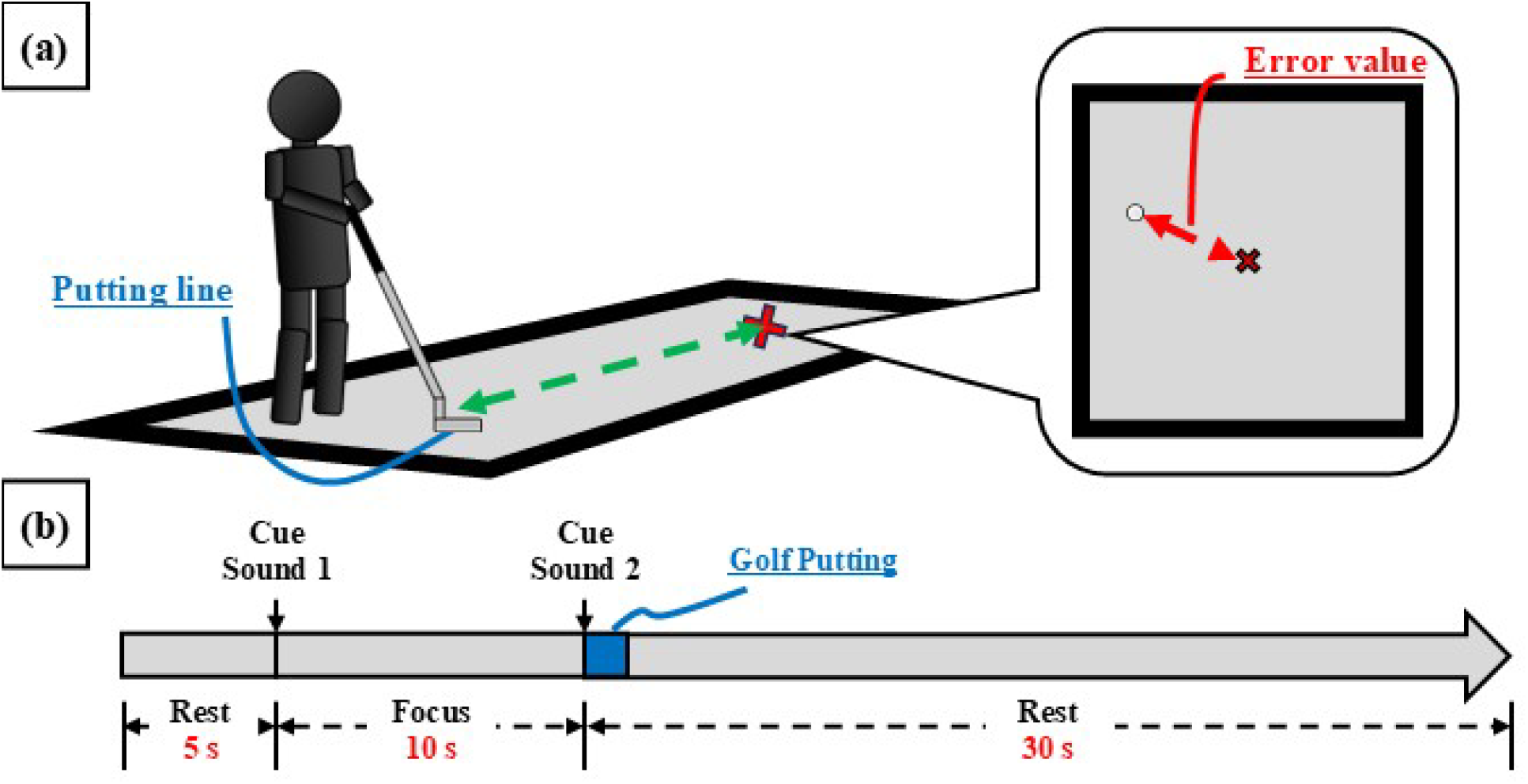
Overview of the putting experiment.

In the competition session, participants performed the putting task under competition-related instructions. Specifically, they were told that prize money could be won based on their putting performance: if their average error value was smaller than that of the experimenter, they would receive 3,000 yen; otherwise, they would receive 1,000 yen. This incentive was chosen with reference to the typical hourly wage for part-time jobs in Japan (approximately 1,000 yen), and a magnitude of two to three times this amount was considered sufficient to induce meaningful psychological pressure. These instructions were designed to elicit psychological pressure through monetary incentives (Hickman & Metz, 2015; Lee & Grafton, 2015). No competition-related instructions were given during the practice session.

The experimental sessions were conducted in the following order: a 3-min skill acquisition session, followed by the practice and competition sessions. The skill acquisition session was included to minimize the potential effects of task habituation on performance.

### fNIRS measurements

A multichannel fNIRS device (WOT-HS, NeU Inc., Japan) was used to measure optical signals associated with hemodynamic changes at a sampling rate of 10 Hz across 56 channels covering the frontal cortex (Atsumori et al., 2009; Burin et al., 2019). Channels 1 to 34 measured optical changes using a light-emitting diode (LED) illuminator (wavelengths of 730 and 850 nm) and a standard-distance detector spaced 30 mm apart. Channels 35–56 used an LED illuminator with the same wavelengths and a short-distance detector spaced 21.2 mm apart. The fNIRS probe set was mounted on the participant’s head according to the international 10-20 system, centered on channel 18 (Fig. 2).

**Figure 2.**
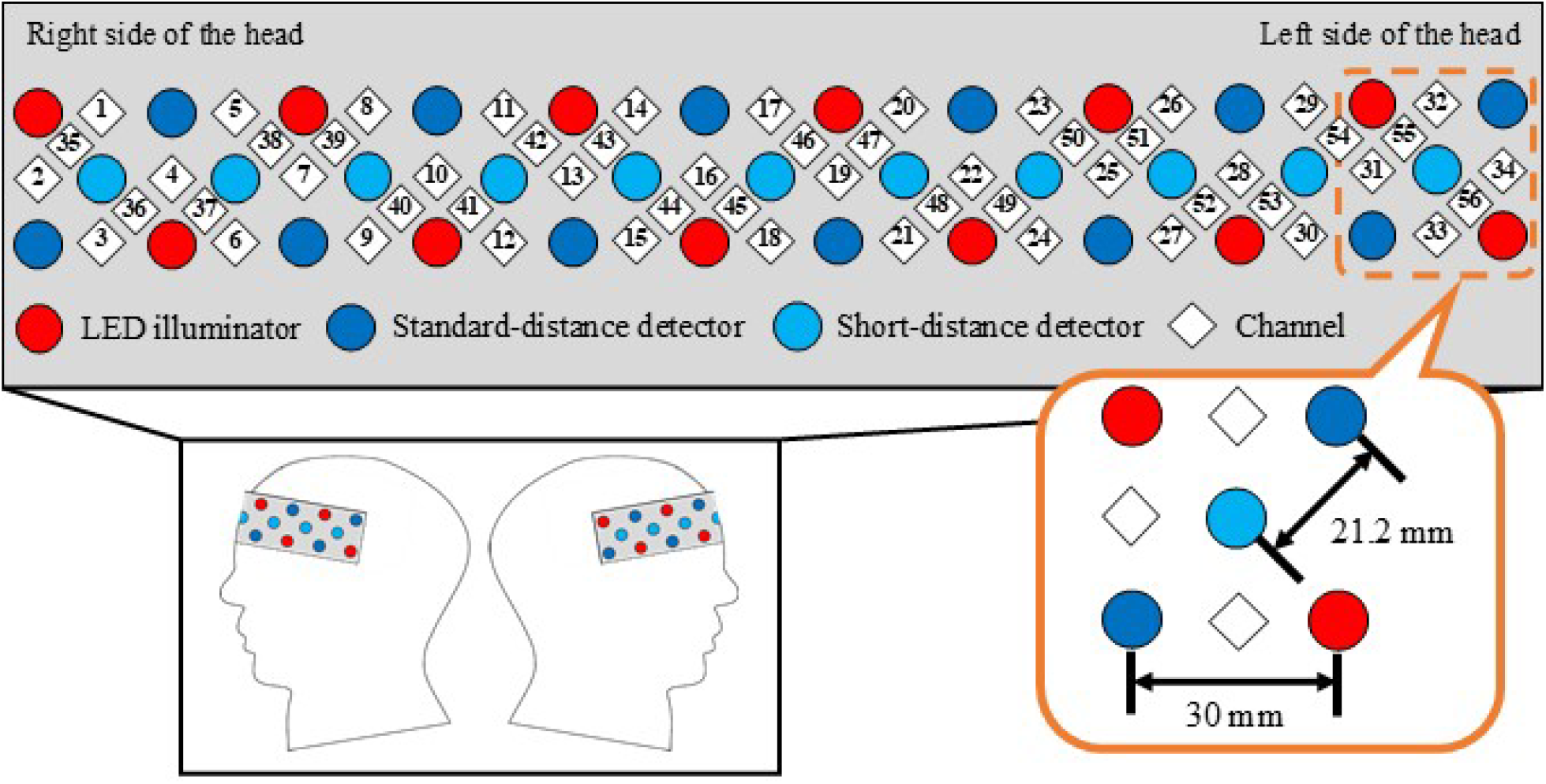
Overview of functional near-infrared spectroscopy device attachment. The light-emitting diode (LED) illuminator is represented by the red circle. The Standard-distance detector is represented by the blue circle. The Short-distance detector is represented by the light blue circle. The channels number are shown in black.

For spatial registration of the fNIRS channel coordinates on the cortical surface, 10 additional participants were measured using a three-dimensional digitizer (EZT-DM401, Hitachi Ltd., Japan). The cortical projection positions were estimated using an fNIRS spatial analysis tool (Okamoto et al., 2004). Corresponding brain regions were estimated based on the spatial coordinate data for each channel using an anatomical labeling tool (Singh et al., 2005). The mean MNI coordinates and anatomical labels for channels 1–34 are shown in Table 1.

**Table 1.**
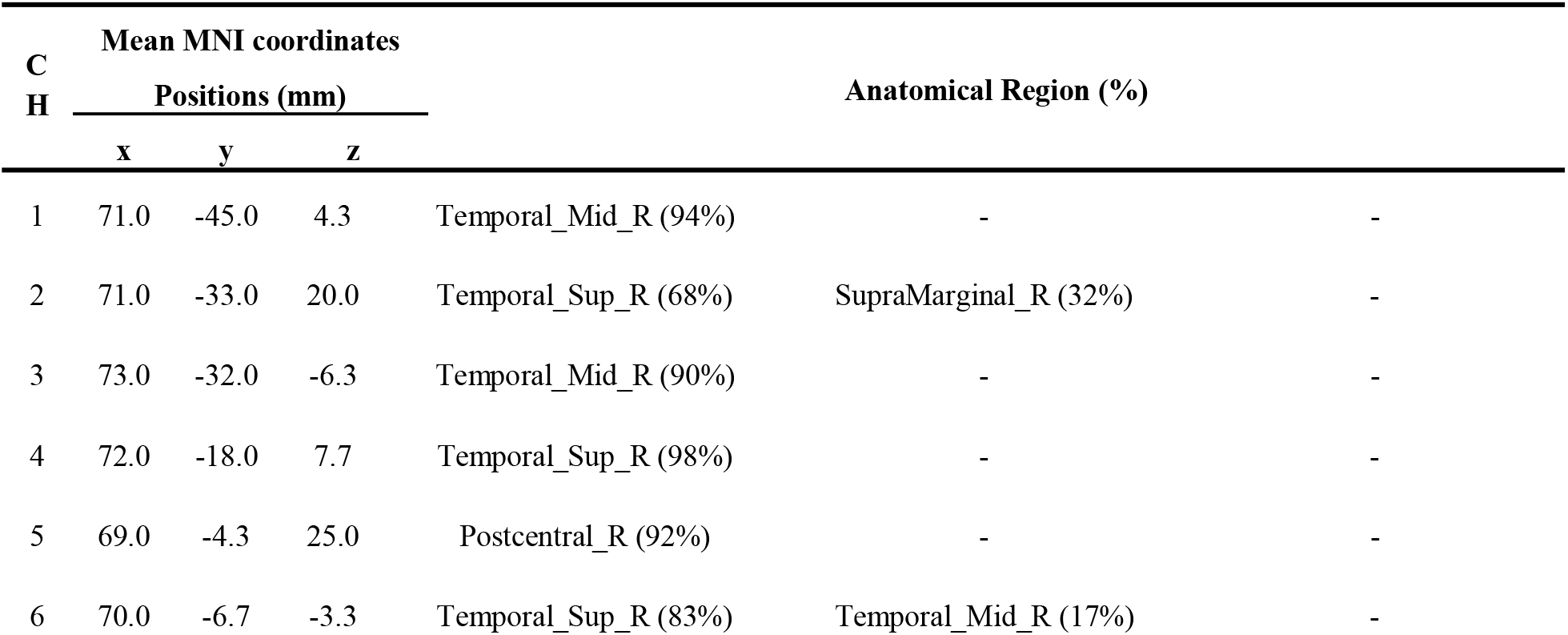
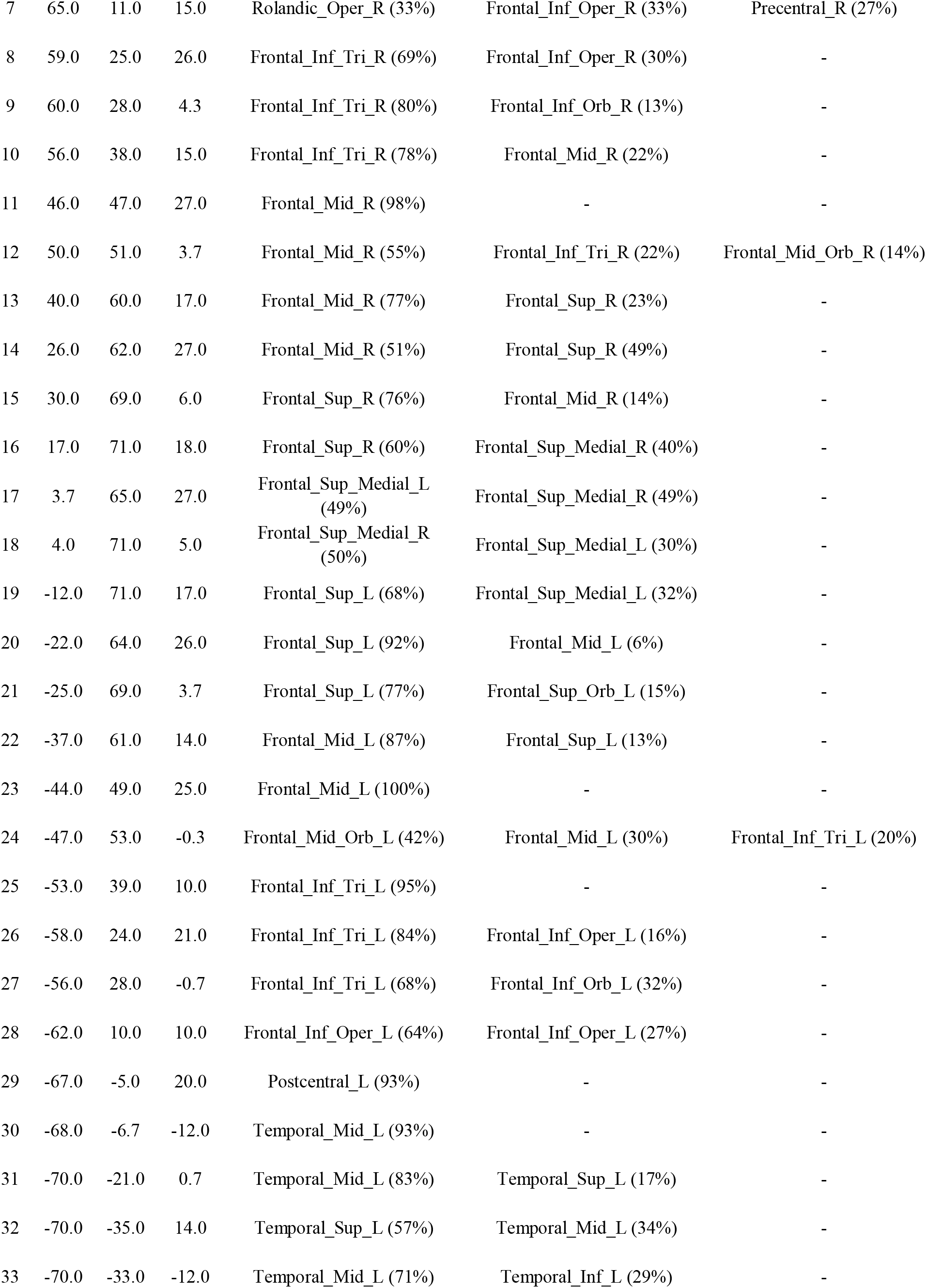
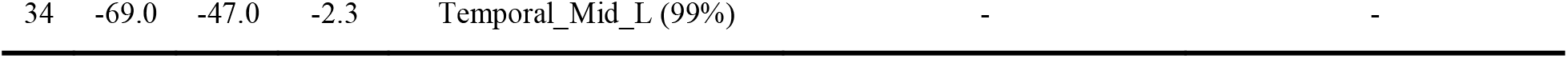
Mean MNI coordinates and anatomical labels for fNIRS measurement channels.

The participants used for the spatial registration were independent from those in the main experiment. The groups were matched in race and age range; therefore, their average spatial coordinates were assumed to be similar.

## Data Analysis

### Classification of participants based on performance

For each participant, the mean error value was calculated separately across the five trials in the practice and competition sessions. Performance change was defined as the Δerror value and calculated by subtracting the mean error value in the practice session from that in the competition session. Accordingly, positive Δerror values indicated performance deterioration, whereas negative Δerror values indicated performance improvement under psychological pressure. Participants with positive Δerror values were classified into the choking group because they failed to maintain their baseline performance under psychological pressure (Mesagno & Hill, 2013). Participants with negative Δerror values were classified into the non-choking group because their performance was not negatively affected by psychological pressure.

### Brain activation and performance

The time series data measured by the fNIRS device were first preprocessed using the Platform for Optical Topography Analysis Tools (POTATo) ver. 3.8.51 (National Institute of Advanced Industrial Science and Technology, Japan) (Sutoko et al., 2016). First, the optical data measured from all 56 channels were converted into hemoglobin (Hb) concentration signals using the modified Beer–Lambert law (Cope & Delpy, 1988). Specifically, these were converted into oxygenated hemoglobin (Oxy-Hb) and deoxygenated hemoglobin (Deoxy-Hb) signals. Second, a third-order Butterworth band-pass filter with a frequency range of 0.01 to 0.1 Hz was applied to the Oxy-Hb and Deoxy-Hb signals in each channel (Naseer & Hong, 2015). Third, the skin blood flow components were removed from channels 1 to 34 using the method proposed by Funane et al. (Funane et al., 2014). This method uses independent component analysis to group channels that include standard-separation (channels 1–34) and short-separation (channels 35–56) Hb signals, to separate the superficial and deep-layer components. The channel combinations used in this analysis are listed in Table 2.

**Table 2.** Channel groups for execution of independent component analysis.

| Channel group no. | Channel (S-D distance = 30.0 [mm]) | Channel (S-D distance = 21.2 [mm]) |
| --- | --- | --- |
| 1 | 1 | 35 |
| 2 | 2 | 35 |
| 3 | 3 | 36 |
| 4 | 4 | 36,37 |
| 5 | 5 | 38 |
| 6 | 6 | 37 |
| 7 | 7 | 38,39 |
| 8 | 8 | 39 |
| 9 | 9 | 40 |
| 10 | 10 | 40,41 |
| 11 | 11 | 42 |
| 12 | 12 | 41 |
| 13 | 13 | 42,43 |
| 14 | 14 | 43 |
| 15 | 15 | 44 |
| 16 | 16 | 44,45 |
| 17 | 17 | 46 |
| 18 | 18 | 45 |
| 19 | 19 | 46,47 |
| 20 | 20 | 47 |
| 21 | 21 | 48 |
| 22 | 22 | 48,49 |
| 23 | 23 | 50 |
| 24 | 24 | 49 |
| 25 | 25 | 50,51 |
| 26 | 26 | 51 |
| 27 | 27 | 52 |
| 28 | 28 | 52,53 |
| 29 | 29 | 54 |
| 30 | 30 | 53 |
| 31 | 31 | 54,55 |
| 32 | 32 | 55 |
| 33 | 33 | 56 |
| 34 | 34 | 56 |

Analysis was performed using the POTATo plug-in developed by the original authors.

Fourth, the Oxy-Hb and Deoxy-Hb signals were segmented into epochs comprising the 10-s focus period before putting for 45 s, including 2 s before and 33 s after the onset of the focus period. Baseline-corrected Oxy-Hb and Deoxy-Hb signals during the focus period were calculated by subtracting the mean of the 2 s pre-focus baseline values from the raw signals. Fifth, putting trials in which baseline-corrected Oxy-Hb or Deoxy-Hb signals showed abrupt changes > 0.2 mM·mm within a 0.2-s interval during the focus period were excluded as motion artifacts (Kawaike et al., 2019). For each participant, the baseline-corrected Oxy-Hb and Deoxy-Hb signals of the remaining valid trials were averaged for the practice and competition sessions. If fewer than three valid trials were available in a given channel for either session, the channel was excluded from further analysis.

Considering the hemodynamic delay relative to neural activity (Dehaene-Lambertz, Dehaene & Hertz-Pannier, 2002), the average Oxy-Hb and Deoxy-Hb amplitudes during the final 7.5 s of the focus period were calculated for both sessions and used as indices of activation (Oxy-Hb and Deoxy-Hb values). Activation changes were quantified by calculating the differences in Oxy-Hb and Deoxy-Hb values between practice and competition sessions and were defined as ΔOxy-Hb and ΔDeoxy-Hb values, respectively.

The Mann–Whitney U test was used to compare ΔOxy-Hb and ΔDeoxy-Hb values between the choking and non-choking groups. For channels showing significant group differences in ΔOxy-Hb values, Spearman’s rank-correlation analysis was performed between the ΔOxy-Hb values and the corresponding Δerror values.

## Results

Among the 20 participants’ fNIRS datasets, three could not be processed owing to preprocessing errors in POTATo (Sutoko et al., 2016). Therefore, the error values and fNIRS data from these participants were excluded from further analyses.

Motion artifact-based trial rejection was applied to the fNIRS datasets of the remaining 17 participants. Consequently, the proportion of valid channels used to compare changes in brain activity was 95.9% (±3.5%) on average, indicating generally high signal quality across participants.

### Classification of participants based on performance

The average error values of participants during the practice and competition sessions are shown in Fig. 3. Of the 17 participants, seven had positive Δerror values, indicating performance deterioration, whereas 10 had negative Δerror values, indicating performance improvement. Accordingly, seven participants were classified into the choking group and 10 into the non-choking group.

**Figure 3.**
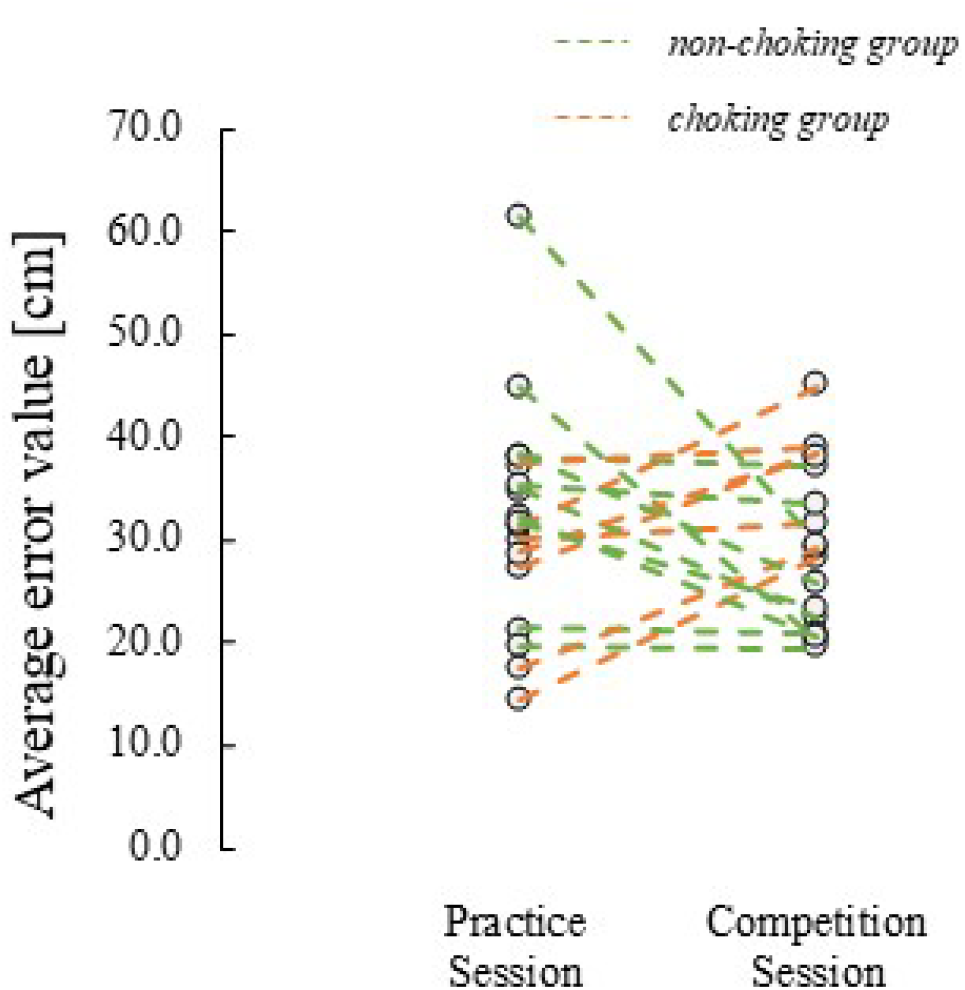
Average error value in practice and competition session.

### Differences in Brain Activation Depending on Performance

The Mann–Whitney U test was performed on ΔOxy-Hb and ΔDeoxy-Hb values for each channel between the two groups to investigate differences in prefrontal cortex (PFC) activation under psychological pressure based on motor performance.

Significant group differences were observed in ΔOxy-Hb values at channels 20 (*U* = 8, *p* = 0.043, *r* = 0.535, uncorrected) and 21 (*U* = 12, *p* = 0.025, *r* = 0.533, uncorrected), with greater increases observed in the non-choking group than in the choking group (Fig. 4).

**Figure 4.**
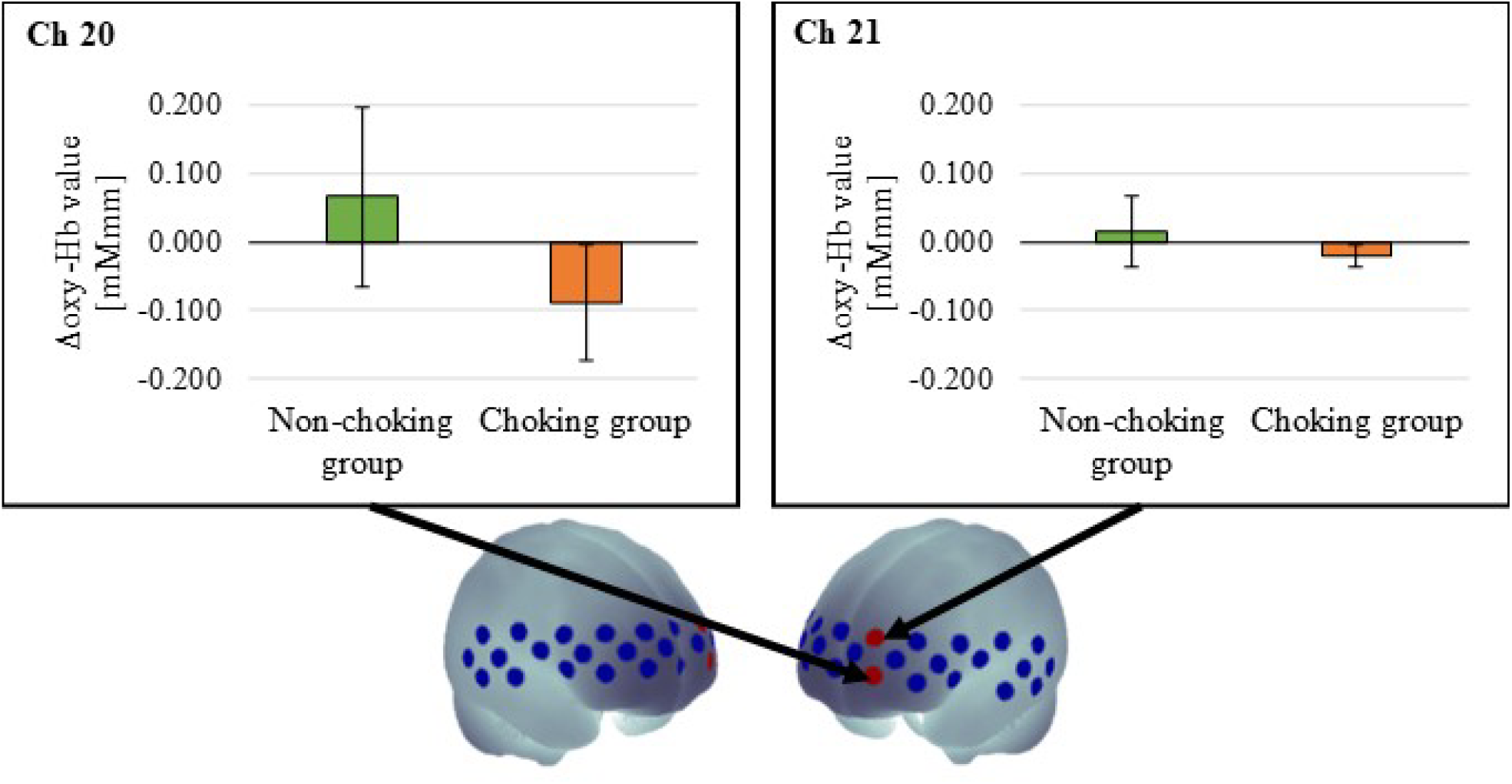
Comparison between the non-choking group and the choking group for change in oxygenated hemoglobin concentration value between practice and competition session. No significant differences were observed in ΔDeoxy-Hb values (channel 20: *U* = 12, *p* = 0.142, *r* = 0.397, uncorrected; channel 21: *U* = 21, *p* = 0.193, *r* = 0.320, uncorrected).

According to the spatial registration results, channels 20 and 21 corresponded approximately to the left superior frontal gyrus. Anatomical labeling identified channels 20 and 21 primarily as Frontal_Sup_L (92% and 77%, respectively; Table 1).

### Correlation Between Brain Activity and Performance change

Spearman’s rank-correlation analysis was performed between ΔOxy-Hb values and Δerror values in channels 20 and 21. No statistically significant correlations were observed. However, moderate negative correlations were observed in channel 20 (ρ = −0.484, *p* = 0.079) and channel 21 (ρ = −0.432, *p* = 0.084) (Fig. 5).

**Figure 5.**
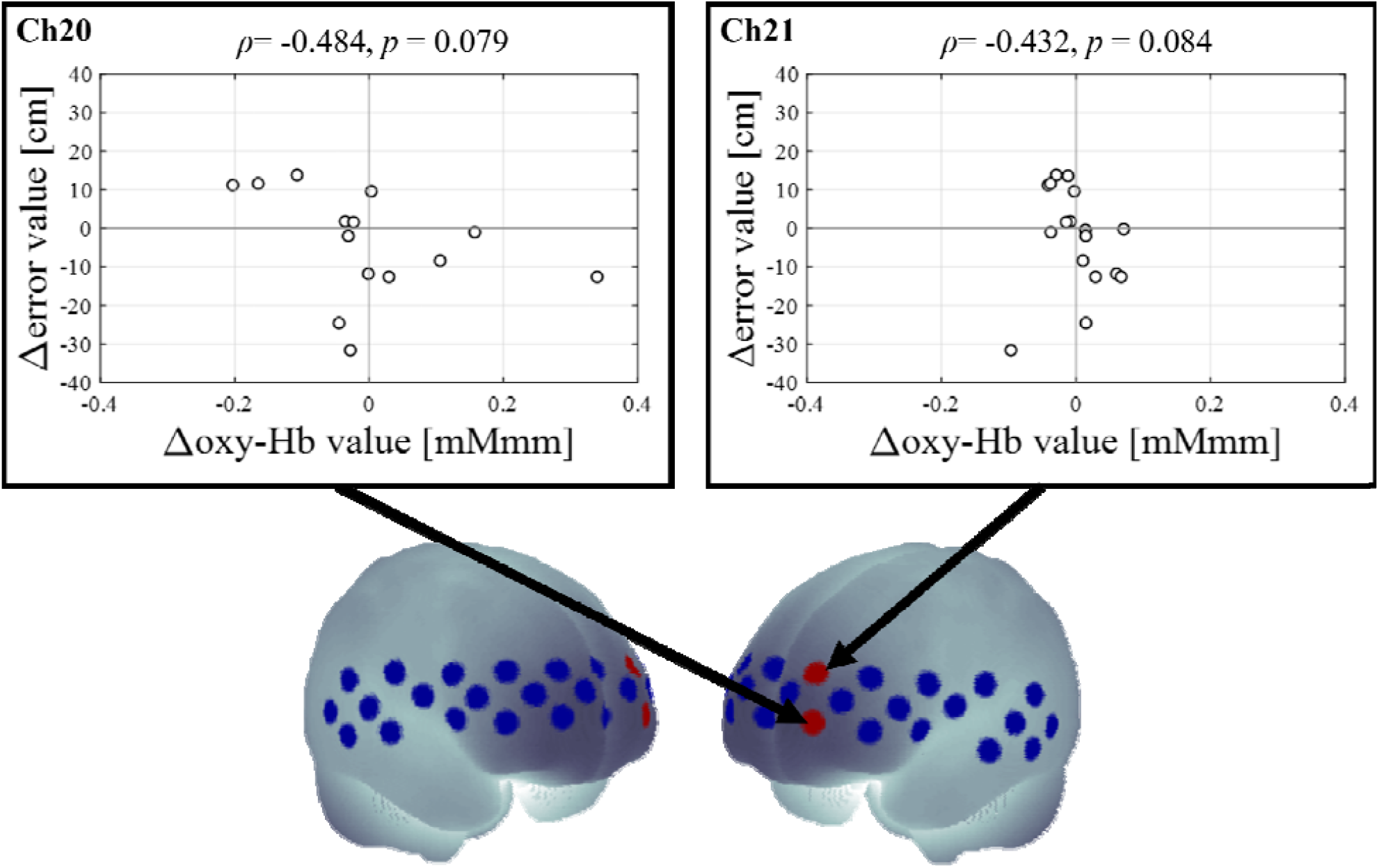
Spearman’s rank correlations between ΔOxy-Hb values and Δerror values in channels 20 and 21.

## Discussion

In this study, we investigated brain activation during golf putting under psychological pressure induced by monetary incentives to develop an NF system that may help individuals overcome choking.

### Inducing Choking

Overall, 17 participants completed five putting attempts in each of the practice and competition sessions. Seven participants had positive Δerror values, indicating performance deterioration under psychological pressure, whereas the remaining 10 participants had negative Δerror values, indicating performance improvement.

Participants performed a 3-min skill acquisition session before the practice and competition sessions to minimize performance variability due to unfamiliarity with the task. Given these experimental conditions, the decline in performance observed during the competition session was attributed to the psychological pressure induced by monetary incentives. These results indicate that choking was successfully induced.

### Performance and brain activity under psychological pressure

We compared the changes in PFC hemodynamic responses between the two groups under psychological pressure using fNIRS. The results showed that ΔOxy-Hb values in the left superior frontal gyrus were significantly greater in the non-choking group than in the choking group.

These findings can be interpreted in light of the distraction theory (Yu, 2015). Specifically, psychological pressure due to monetary incentives may have led to task-irrelevant thoughts such as “*What if I fail?*” or “*I must not fail*.” This distraction possibly prevented participants from allocating sufficient cognitive resources to the putting task, ultimately leading to a decline in performance.

From a neurophysiological perspective, previous studies have shown a close association between the activity in the left superior frontal gyrus and working memory (Cutini et al., 2008; Furley & Wood, 2016; Alagapan et al., 2019). In this study, significant between-group differences in ΔOxy-Hb values were observed at channels 20 and 21, both of which corresponded to this region. These findings suggest that psychological pressure may suppress activity in the left superior frontal gyrus, impairing working memory (reducing cognitive resources owing to increased task-irrelevant thoughts), which may lead to decreased performance.

Moreover, the difference between participants who maintained their performance level and those who showed declined levels may be attributable to differences in their working memory capacity. Previous studies have suggested that working memory capacity can predict performance under pressure (Wood, Vine & Wilson, 2016). Based on these findings, we propose that an NF system targeting the left superior frontal gyrus can enhance working memory and help individuals overcome choking.

### Correlation between brain activity and performance changes

We further examined the correlations between ΔOxy-Hb values and Δerror values in channels 20 and 21. Although neither correlation reached statistical significance, moderate negative correlations were observed in channel 20 (ρ = −0.484, *p* = 0.079) and channel 21 (ρ = −0.432, *p* = 0.084). Because positive Δerror values indicated performance deterioration, these negative correlations suggest that greater increases in ΔOxy-Hb were associated with less performance deterioration or greater performance improvement under psychological pressure. Taken together with the group-comparison results, these findings suggest that activation changes in the left superior frontal gyrus warrant further investigation as a candidate biomarker for NF interventions aimed at mitigating choking under pressure. However, given the exploratory design and limited sample size, these correlations should be interpreted cautiously.

## Limitations

This study had some limitations. First, whether the experimental design accurately reproduces real competitive situations, where psychological pressure affects motor performance, is unclear. In actual competitions, athletes often face situations where the outcome depends on a single attempt. In these cases, psychological pressure is expected to significantly affect performance.

For instance, in golf, making or missing a decisive putt in the 18th hole can determine victory or defeat. Similarly, in soccer, a penalty shootout after extra time, where one successful or missed shot decides the match, illustrates the critical impact of psychological pressure. However, in this study, we prioritized the robustness of the fNIRS experimental design and used a block approach (Luke et al., 2021). This approach may limit the ecological validity; nonetheless, it enhances the reliability of fNIRS measurements. Future studies should refine the experimental and analytical designs to better replicate real competitive conditions.

Second, the statistical analyses were performed without applying multiple comparison correction. When Bonferroni correction (Fishburn et al., 2014), false discovery rate correction (Singh & Dan, 2006), and permutation tests (Pellegrino et al., 2016) were applied, statistical significance was lost in all cases. Therefore, we cannot dismiss the possibility that these results were obtained by chance. Nevertheless, hemodynamic changes associated with neural activity often occur in spatially extended regions of the brain (Pereira et al., 2023). the consistent trends observed in channels 20 and 21, where fNIRS activity was greater in the group with increased activation than in the group with decreased activation, may have some validity owing to their spatial proximity. Additionally, the sample size in this study was determined heuristically, which may have been insufficient. Therefore, future studies should determine the appropriate sample size quantitatively and verify the reproducibility of these findings.

Finally, our findings were derived from exploratory rather than preregistered analyses. Exploratory analyses are valuable for generating new hypotheses; however, they carry the risk of producing chance findings, which can reduce the statistical reliability and generalizability of the results (Wagenmakers et al., 2012). To enhance reproducibility, analysis methods should be predefined, and discretion in data processing should be limited (Nosek et al., 2018). Therefore, this study should be considered as a feasibility study. In future research, we intend to adopt preregistration and registered reports and conduct replication experiments using predefined analysis pipelines to determine the robustness and reliability of the findings (van ‘t Veer & Giner-Sorolla, 2016).

## Conclusion

In this study, we conducted an experiment in which participants performed a golf-putting task under psychological pressure induced by monetary incentives. The aim was to explore neural features that could inform the development of an NF system to help overcome choking. Using fNIRS, we investigated hemodynamic features associated with task performance. Based on Δerror values, participants were categorized into the choking and non-choking groups, and frontal cortex activation was compared between the two groups. The results showed that activation of the left superior frontal gyrus was significantly greater in the non-choking group than in the choking group. In addition, moderate but statistically nonsignificant negative correlations were observed between ΔOxy-Hb values and Δerror values in channels 20 and 21.

These exploratory findings suggest that greater activation increases in the left superior frontal gyrus may be associated with better maintenance or improvement of golf-putting performance under psychological pressure. Therefore, this region warrants further investigation as a candidate biomarker for NF interventions aimed at mitigating choking.

Based on these results, we plan to quantitatively determine the required sample size and examine the reproducibility of our findings in future studies. Furthermore, we aim to design an NF system informed by these results and investigate whether NF training could help individuals overcome choking under pressure.

## Acknowledgements

We thank Mr. Hideyuki Oshima, Mr. Tatsuru Ueda, and Mr. Kodai Takano for their technical assistance. We also thank Prof. Ryota Akagi for the meaningful discussions.

We used ChatGPT 5.1 solely to refine grammar and improve the clarity of English expressions during the preparation of this manuscript. We also thank Editage (www.editage.com) for providing English language editing services.

## Notes

### Competing Interest Statement

The authors have declared no competing interest.

### Funding

- Japan Society for the Promotion of Science: JP20K11367

